# Class A PBPs have a distinct and unique role in the construction of the pneumococcal cell wall

**DOI:** 10.1101/665463

**Authors:** Daniel Straume, Katarzyna Wiaroslawa Piechowiak, Silje Olsen, Gro Anita Stamsås, Kari Helene Berg, Morten Kjos, Maria Victoria Heggenhougen, Martin Alcorlo, Juan A. Hermoso, Leiv Sigve Håvarstein

## Abstract

In oval shaped *Streptococcus pneumoniae*, septal and longitudinal peptidoglycan synthesis is performed by independent functional complexes; the divisome and the elongasome. Penicillin binding proteins (PBPs) were long considered as the key peptidoglycan synthesizing enzymes in these complexes. Among these were the bifunctional class A PBPs, which are both glycosyltransferases and transpeptidases, and monofunctional class B PBPs with only transpeptidase activity. Recently, however, it was established that the monofunctional class B PBPs work together with non-PBP glycosyltransferases (FtsW and RodA) to make up the core peptidoglycan synthesizing machineries within the pneumococcal divisome (FtsW/PBP2x) and elongasome (RodA/PBP2b). The function of class A PBPs is therefore now an open question. Here we utilize the peptidoglycan hydrolase CbpD to show that class A PBPs have an autonomous role during cell wall synthesis in *S. pneumoniae*. Purified CbpD was shown to target the septum of *S. pneumoniae* cells. Using assays to specifically inhibit PBP2x, we demonstrate that CbpD specifically target nascent peptidoglycan synthesized by the divisome. Notably, class A PBPs could process this nascent peptidoglycan from a CbpD-sensitive to a CbpD-resistant form. The class A PBP mediated processing was independent of divisome and elongasome activities. Class A PBPs thus constitute an autonomous functional entity which processes or repairs nascent peptidoglycan synthesized by FtsW/PBP2x. Our results support a model in which pneumococcal peptidoglycan is made by three functional entities, the divisome, the elongasome and a peptidoglycan-repairing or -remodelling complex consisting of bifunctional PBPs. To our knowledge this is the first time a specific function has been identified for class A PBPs in bacterial cell wall synthesis.

## Introduction

The peptidoglycan layer covering the pneumococcal cell provides shape and rigidity, and is essential for growth and survival. It consists of linear chains of two alternating amino sugars, N-acetylglucosamine (GlcNAc) and N-acetylmuramic acid (MurNAc), interlinked by peptide bridges between MurNAcs on adjacent strands^1,2^. Peptidoglycan is synthesized from lipid II precursors at the outside the cytoplasmic membrane by glycosyltransferases that polymerize the glycan chains and transpeptidases that interconnect the chains through peptide cross-links. *S. pneumoniae* produce five different <u>p</u>enicillin-<u>b</u>inding <u>p</u>roteins (PBPs) with transpeptidase activity, namely PBP1a, PBP1b, PBP2a, PBP2b and PBP2x^3^. The first three of these, designated class A PBPs, are bifunctional enzymes that catalyse transglycosylation as well as transpeptidation, while PBP2x and PBP2b are monofunctional transpeptidases (class B PBPs)^4^. Monofunctional glycosyltransferases that have homology to the glycosyltransferase domains of class A PBPs are present in some bacterial species, but are absent from *S. pneumoniae*. PBP2x is an essential constituent of the divisome, a multiprotein division machine that synthesizes the septal cross-wall^3,5,6,7^. The other monofunctional transpeptidase, PBP2b, is a key component of another multiprotein complex, the elongasome, which is responsible for longitudinal peptidoglycan synthesis^3,5,6,7,8^. Until recently, it was believed that only class A PBPs were able to polymerize glycan chains in *S. pneumoniae*. Consequently, the divisome as well as the elongasome would have to include at least one class A PBP in order to be functional. Recently, however, it was discovered that FtsW and RodA, two proteins belonging to the SEDS (<u>s</u>hape, <u>e</u>longation, <u>d</u>ivision, and <u>s</u>porulation) family, function as peptidoglycan polymerases that synthesize glycan strands from lipid II^9,10,11^. FtsW and RodA were originally reported to be lipid II flippases, a function now assigned to MurJ^12^. However, it is still not entirely clear whether these polytopic membrane proteins are monofunctional glycan polymerases or bifunctional flippases and polymerases^13,14^. Previous research has shown that FtsW and RodA are essential, and work in conjunction with PBP2x and PBP2b, respectively^9,11^.

Peptidoglycan synthesis requires the concerted action of enzymes that carry out transglycosylation and transpeptidation reactions. Thus, in principle, peptidoglycan synthesis might be performed by monofunctional transglycosylase working together with monofunctional transpeptidase, by single bifunctional enzymes such as the class A PBPs, or by a combination of monofunctional and bifunctional enzymes. As mentioned above, class A PBPs have traditionally been considered to be essential components of bacterial divisomes and elongasomes. However, it has been known for a long time that *Bacillus subtilis* is viable without class A PBPs^15^. Thus, considering the recent discovery of the SEDS partners of PBP2x and PBP2b, it is conceivable that the pneumococcal divisome and elongasome perform the primary synthesis of septal and peripheral peptidoglycan without the involvement of class A PBPs. If so, the function of class A PBPs is an open question, and their role in peptidoglycan synthesis must be re-examined. Here, we have addressed this question by exploiting the unique properties of the peptidoglycan hydrolase CbpD (<u>c</u>holine-<u>b</u>inding <u>p</u>rotein <u>D</u>).

When developing competence for natural transformation, streptococci belonging to the mitis phylogenetic group express a set of core competence genes controlled by the competence stimulating peptide (CSP)^16^ and the competence-specific two-component regulatory system ComDE^17^. Most proteins whose expression are controlled by this quorum-sensing-like system are involved in DNA-binding, DNA-uptake, DNA-processing or genomic integration of internalized DNA^18,19^. However, among the CSP/ComDE-regulated genes is a peptidoglycan-degrading enzyme, CbpD, which does not seem to have a role in any of the transformation steps mentioned above^19,20^. Instead, evidence strongly suggests that CbpD, which is encoded by a late competence gene, is part of a DNA-acquisition mechanism consisting of CbpD and the cognate immunity protein ComM^21,22,23^. Previous research has shown that ComM, a polytopic membrane protein encoded by an early competence gene, protects competent pneumococci from being lysed by their own CbpD. The mechanism behind this protection is not understood^24,25^.

CbpD is composed of three different domain types: an N-terminal <u>c</u>ysteine, <u>h</u>istidine-dependent <u>a</u>midohydrolase/<u>p</u>eptidase (CHAP) domain, one or two Src homology 3b (SH3b) domains, and a C-terminal choline-binding domain (Cbd) consisting of four choline-binding repeats. CHAP domains are present in many peptidoglycan hydrolases, and function as either N-acetylmuramoyl-L-alanine amidases or endopeptidases^19,26^. Hence, the CHAP domain of CbpD must cleave somewhere within the peptide bridges of streptococcal peptidoglycan. However, so far the exact bond cleaved has not been identified. The SH3b domain is essential for the function of CbpD, and experimental evidence indicates that it binds to the peptidoglycan portion of the cell wall^27^. CbpDs from *Streptococcus mitis* and *Streptococcus oralis* contain only one SH3b domain, sandwiched between the CHAP and the Cbd domain, while many (but not all) strains of *S. pneumoniae* contain two successive SH3b domains. The choline-binding repeats of the Cbd domain anchor CbpD to cell wall teichoic acid, and possibly also lipoteichoic acid, through non-covalent interactions with the choline residues decorating these polymers^28^. Mitis group streptococci such as *S. pneumoniae, S. pseudopneumoniae, S. mitis, S. oralis* and *S. infantis* produce choline-decorated teichoic acids, while this type of teichoic acid is not present in mitis group streptococci that are more distantly related to *S. pneumoniae*^29^. Similar to the CHAP and SH3b domains, the Cbd domain is essential for the biological function of CbpD^27^.

Even though CbpD appears to be a key component of the pneumococcal gene transfer machinery it is still poorly characterized. It has proved very difficult to express pneumococcal CbpD in a soluble and active form in *Escherichia coli* and other hosts, and we therefore investigated whether homologs of pneumococcal CbpD from various mitis group streptococci might be more amenable to heterologous expression. This strategy turned out to be successful, as we were able to purify milligram amounts of the CbpD protein produced by *S. mitis* B6 (CbpD-B6) in soluble form. In the present study, we show that CbpD-B6 is active against *S. pneumoniae*, and that its properties appear to be very similar to pneumococcal CbpD. Interestingly, we found that CbpD-B6 cleaves only a distinct subset of the peptide bridges that cross-link the carbohydrate chains in pneumococcal peptidoglycan. It is highly specific for nascent peptidoglycan formed by PBP2x and FtsW. We realized that this property can be exploited as a research tool. Hence, we have used the unique specificity of CbpD to study the functional relationships between different peptidoglycan synthesizing enzymes in *S. pneumoniae*. Our results strongly indicate that class A PBPs are not part of the core machinery of the divisome and elongasome, but have an important autonomous role in construction of the fully matured peptidoglycan layer.

## Results

### Purification and properties of CbpD-B6

The gene encoding *cbpD* from *S. mitis* B6 was amplified by PCR, ligated into the pRSET-A vector, and expressed using *E. coli* BL21 cells as a host. Since choline-binding proteins bind to the diethylaminoethanol groups of DEAE-cellulose via their choline-binding repeats, the soluble protein fraction was subjected to DEAE-cellulose affinity chromatography as previously described^30^. The recombinant CbpD-B6 protein was further purified by size-exclusion chromatography (SEC) on a Superdex^™^ 75 10/300 GL column (see Methods section for details). After SEC some CbpD-B6 fractions were essentially pure as determined by SDS-PAGE (Fig. S1). These fractions were collected and used for further studies.

CbpD-B6 and CbpD from *S. pneumoniae* strain R6 (CbpD-R6) are highly homologous except for the extra SH3b domain present in CbpD-R6 (Fig. S2). We therefore assumed that the two enzymes have more or less the same properties, and that CbpD-B6 can be used as a proxy for CbpD-R6. To test this assumption, we first investigated whether CbpD-B6 is active against non-competent *S. pneumoniae* R6 cells. The results presented in Fig. 1A show that the R6 strain is highly sensitive to CbpD-B6, and that a concentration of about 0.3 µg ml^−1^ lyses 50% of the cells in a R6 culture at OD_550_ = 0.2. To compare the properties of the CbpD-R6 and CbpD-B6 enzymes under *in vivo* conditions, we replaced the native *cbpD* gene with *cbpD-B6* in strain R6. This was done in a wild-type strain (A) as well as a Δ*comM* genetic background (strain B), resulting in strains C (strain A, *cbpD-*B6) and D (strain B, *cbpD*-B6). Induction of competence in cultures of these strains by addition of CSP at OD_550_ = 0.2 resulted in almost identical growth and lysis patterns (Fig. 1B and C). The growth of the A and C strains were virtually unaffected by competence induced CbpD secretion, as they express the immunity protein ComM. However, significant and similar lysis were observed when strains B and D were induced to competence (Fig. 1C). These results show that CbpD-R6 and CbpD-B6 have very similar properties.

**Fig. 1.**
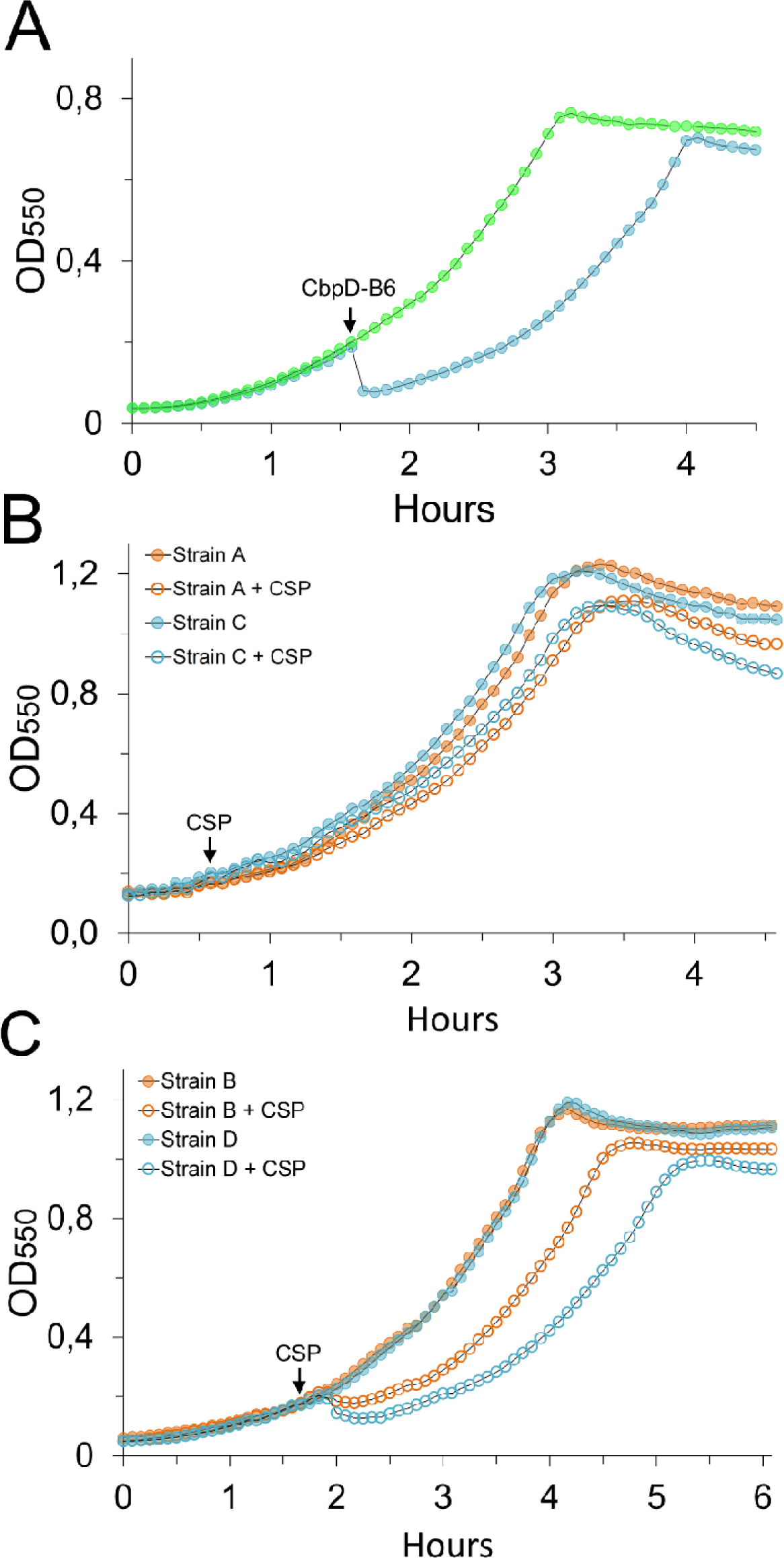
Functionality of CbpD-B6 in *S. pneumoniae*. A. Exponential growing pneumococci (strain RH425) was added purified CbpD-B6 (arrow) to a final concentration of 0.3 µg ml^−1^ (blue circles). Growth of non-treated cells are represented by green circle. B and C. Wild type *S. pneumoniae* (strain A), pneumococci expressing CbpD-B6 from the native *cbpD*-locus (strain C) and their respective Δ*comM* mutants (strain B and D) were induced to competence by adding CSP to a final concentration of 250 ng ml^−1^ (arrows).

### CbpD-B6 attacks the septal area of the pneumococcal cell wall

In an attempt to identify the specific bond cleaved by CbpD, purified peptidoglycan from *S. pneumoniae* R6 was digested with CbpD-B6 and mutanolysin followed by HPLC analysis of the resulting digest^31^. We were not able to detect any significant difference in muropeptide composition between mutanolysin digested samples and CbpD-B6/mutanolysin double-digested samples. This indicated that the enzyme is not active against purified peptidoglycan, or that it only cleaves a small subset of the peptide bonds present in pneumococcal peptidoglycan. To investigate the latter possibility, pneumococci exposed to purified recombinant CbpD-B6 were examined by scanning (SEM) and transmission electron microscopy (TEM) for visualization of changes in their ultrastructure. Pneumococcal cultures were treated for 60 seconds with 0.5 µg ml^−1^ of enzyme, after which the bacteria were fixed and the reaction stopped by the addition of 2% formaldehyde and 2.5% glutaraldehyde (v v^−1^). The SEM microscopy analysis clearly showed that CbpD-B6 attacks only the septal region of the peptidoglycan sacculus, resulting in cells that are split in half along their equators (Fig. 2A and B). Interestingly, the rims of both hemispheres in the split cells are thicker than the rest of the peptidoglycan layer. This suggests that CbpD-B6 cleaves the cells along the middle of the equatorial ring, also called the piecrust. Alternatively, the peptidoglycan layer has rolled back on itself to become double at the rims. In accordance with the SEM data, the TEM micrographs of CbpD-B6 treated cells showed damage to the peptidoglycan layer only in the septal region. In some cells that were about to rupture, “footprints” of the enzyme’s activity could be seen. Judging from these footprints, CbpD-B6 cleaves peptide bridges located within and/or close to the base of the septal cross-wall (Fig. 2C and D).

**Fig. 2.**
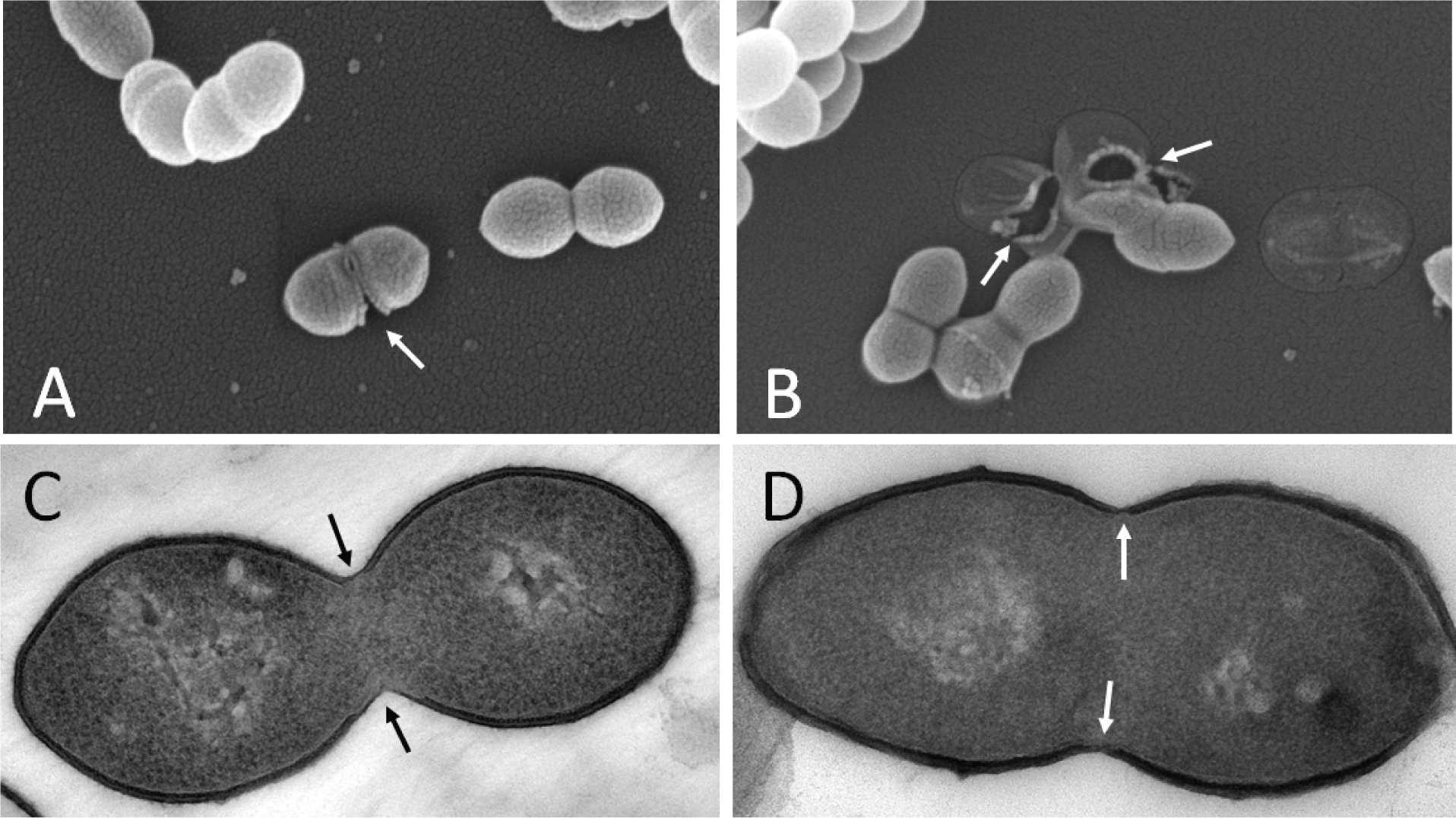
Scanning (A and B) and transmission (C and D) electron micrographs of pneumococci subjected to 0.5 µg ml^−1^ CbpD-B6 for 60 seconds before they were fixed and prepared for electron microscopy. Arrows indicate areas in the cell wall attacked by the muralytic enzyme.

### CbpD-B6 specifically cleaves nascent peptidoglycan formed by PBP2x and FtsW

Since CbpD-B6 attacks the septal region of the cell, we speculated that the enzyme targets the peptidoglycan formed by PBP2x and FtsW. If so, specific inhibition of the divisome activity might render pneumococci less sensitive or insensitive to CbpD-B6. In a recent profiling of the β-lactam selectivity of pneumococcal PBPs, Kocaoglu *et al*.^32^ showed that PBP2x is more sensitive than PBP1a, PBP1b, PBP2a and PBP2b to several different β-lactams. Hence, by using the appropriate β-lactam at the right concentration it should be possible to inhibit the transpeptidase activity of PBP2x without significantly affecting the function of the other PBPs. To test this hypothesis, we grew pneumococcal cultures in 96 well plates in a microplate reader at 37 °C. When reaching OD_550_ ∼ 0.2, each culture was treated with a different concentration of oxacillin. The oxacillin concentrations used ranged from 0-100 µg ml^−1^, i.e. from sub- to supra-MIC concentrations. Ten minutes after being exposed to oxacillin, each culture received 5 µg ml^−1^ of purified CbpD-B6. Comparison of the lytic responses of the cultures showed that the extent of lysis gradually decreased with increasing oxacillin concentrations until the cells became resistant to CbpD-B6 at concentrations between 0.19 – 6.1 µg ml^−1^ (Fig. 3A). The lowest concentration that gave full protection (0.19 µg ml^−1^), corresponds roughly to the MIC value of oxacillin against the R6 strain (Fig. S3). However, to our great surprise, the pneumococci started to lyse again when the concentration of oxacillin was increased further, i.e. above 6.1 µg ml^−1^. At the highest oxacillin concentrations used (50 and 100 µg ml^−1^), the pneumococci became as sensitive as untreated control cells (Fig. 3A). In sum, the results show that as the oxacillin concentration is gradually increased the lytic response to CbpD-B6 shifts from decreasing sensitivity (S1-phase) to resistance (R-phase) and then back to increasing sensitivity (S2-phase).

**Fig. 3.**
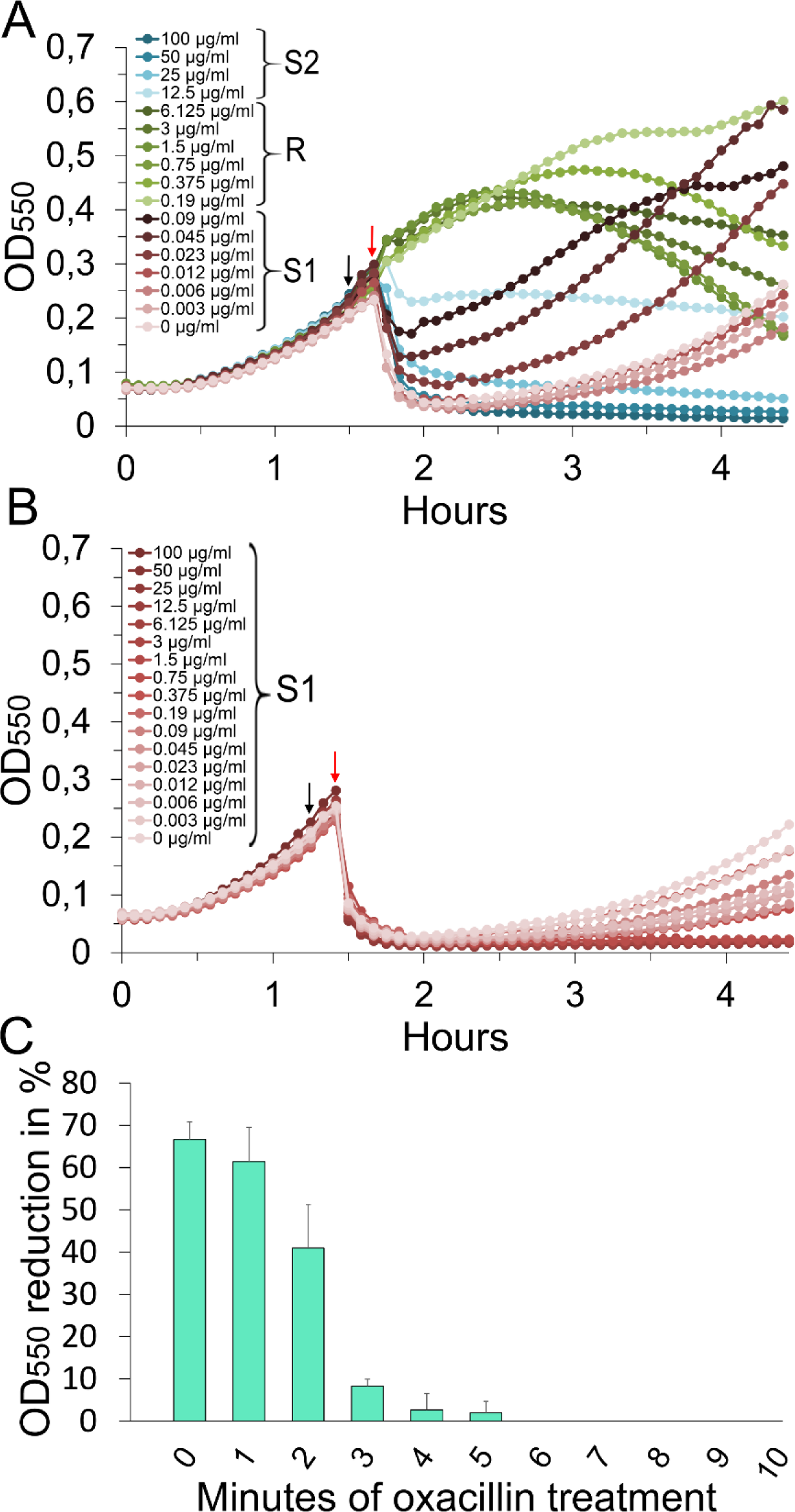
Inhibition of PBP2x results in CbpD-resistant peptidoglycan. A. Increasing concentrations of oxacillin was added to exponentially growing wild type cells (RH425) at OD_550_≈ 0.25 (black arrow). After ten minutes, CbpD-B6 was added (red arrow) to a final concentration of 5 µg ml^−1^. The cells were susceptible to CbpD-B6 from 0-0.09 µg ml^−1^ of oxacillin (S1 phase, red curves), resistant from 0.19-6.125 µg ml^−1^ (R phase, green curves) and susceptible from 12.5-100 µg ml^−1^ (S2 phase, blue curves). B. Oxacillin treatment of pneumococci expressing a low affinity PBP2x (strain KHB321) did not result in CbpD-resistance, but arrested the cells in the S1 phase. C. Wild type cells at OD_550_ ≈ 0.25 were treated with oxacillin (0.8 µg ml^−1^) for 0-10 minutes before CbpD-B6 (5 µg ml^−1^) was added. The percent reduction in cell density caused by cell lysis was determined. Full resistance against CbpD-B6 was developed after about 4-6 minutes of oxacillin treatment. The experiments were repeated three times with the same results.

In line with the observations above, GFP-CbpD has previously been shown to mainly bind the septal region of pneumococcal cells, and the binding specificity is determined by the C-terminal choline-binding domain^27^. To test whether CbpD-resistance during the R-phase could be explained by altered binding of CbpD after exposure to oxacillin, we analyzed the binding patterns of sfGFP-CbpD-B6 as previously described^27^. The fusion protein was expressed and purified essentially as CbpD-B6, and exposed to RH425 control cells as well as RH425 cells treated with 0.8 µg/ml oxacillin for 10 minutes (resulting in R-phase cells, Fig. 3A). sfGFP-CbpD-B6 retained the localization to the septal region after oxacillin-treatment for cells in all division stages (Fig. S4A), although the fraction of cells without septal sfGFP-CbpD-B6 was slightly higher than in the control cells (6.7 % in control cells and 11.8 % after oxacillin treatment, Fig. S4B). This show that the R-phase cannot be explained by alterations in the binding pattern of sfGFP-CbpD-B6.

Beta-lactam-resistant pneumococci have acquired so-called low-affinity PBPs, modified PBPs that have much lower affinity for β-lactams than the corresponding PBPs of sensitive strains. To verify that the R-phase is due to inhibition of PBP2x by oxacillin, the experiment described above was repeated with an R6 mutant strain (KHB321) expressing a low-affinity version of PBP2x. The KHB321 mutant was constructed by replacing the extracytoplasmic part of R6-*pbp2x* with the corresponding part of the low-affinity *pbp2x* gene from *S. mitis* strain B6. The B6 strain is a highly penicillin-resistant clinical isolate that produces low-affinity versions of PBP2x, PBP2b and PBP1a^33^. When the oxacillin titration experiment was carried out with the KHB321 strain, no R-phase was obtained within the concentration range used (0-100 µg ml^−1^ oxacillin) (Fig. 3B). This result clearly shows that inhibition of the transpeptidase activity of PBP2x by oxacillin causes the R-phase. Moreover, it shows that CbpD-B6 specifically attacks the peptidoglycan formed by PBP2x/FtsW in the divisome.

### The S2 phase

During the S1-phases the oxacillin concentration increases gradually resulting in progressively stronger inhibition of PBP2x. This causes a gradual reduction in the number of nascent peptide bridges formed by PBP2x, and eventually complete inhibition of its transpeptidase activity in the R-phase. While this line of reasoning provides an explanation for the S1- and R-phases, it does not explain the S2-phase. How can a further increase in oxacillin concentration lead to increased cell lysis when PBP2x is completely inhibited? We observed that the R-phase disappears if oxacillin (0.8 µg ml^−1^) and CbpD-B6 are added simultaneously to pneumococcal cultures. After being exposed to oxacillin it takes 3-4 minutes before the cells develop full immunity against CbpD-B6 (Fig. 3C). This shows that the peptidoglycan initially formed by the PBP2x/FtsW machinery must undergo remodelling before it becomes resistant to CbpD-B6, a process that takes several minutes. This finding suggested a plausible explanation for the S2-phase. Although PBP2x is more sensitive than the other pneumococcal PBPs to oxacillin, a further increase in oxacillin concentration will eventually affect the transpeptidase activity of the less sensitive PBPs. Presumably, the activity of one or more of these PBPs is required to remodel PBP2x/FtsW-synthesized peptidoglycan into a CbpD-B6-resistant form. Consequently, the cells will not become resistant if their activity is blocked. The reason for this is that newly synthesized CbpD-B6-sensitive peptidoglycan will still be present if the activities of PBP2x and the PBP(s) required for processing this peptidoglycan are blocked simultaneously. In sum, our results indicate that the S2-phase results from inhibition of the PBP(s) required for processing PBP2x/FtsW-synthesized peptidoglycan into a CbpD-B6-resistant form.

### Peptidoglycan synthesized by the FtsW/PBP2x machinery is further processed by class A PBPs

To determine whether class A PBPs are required to produce mature CbpD-B6-resitant peptidoglycan, the oxacillin titration experiment described above was performed in the presence of 10 µg ml^−1^ of the antibiotic moenomycin. Moenomycin inhibits bacterial growth by blocking the transglycosylase activity of class A PBPs, but does not affect FtsW and RodA^9^. Our results showed that in the presence of moenomycin the S1-R-S2 pattern disappeared, and the pneumococci were sensitive to CbpD-B6 at all oxacillin concentrations used (0-100 µg ml^−1^ oxacillin) (Fig. 4 and supplementary Fig. S5A). This demonstrates that without functional class A PBPs, nascent peptidoglycan is not converted to the CbpD-B6-resistant form.

**Fig. 4.**
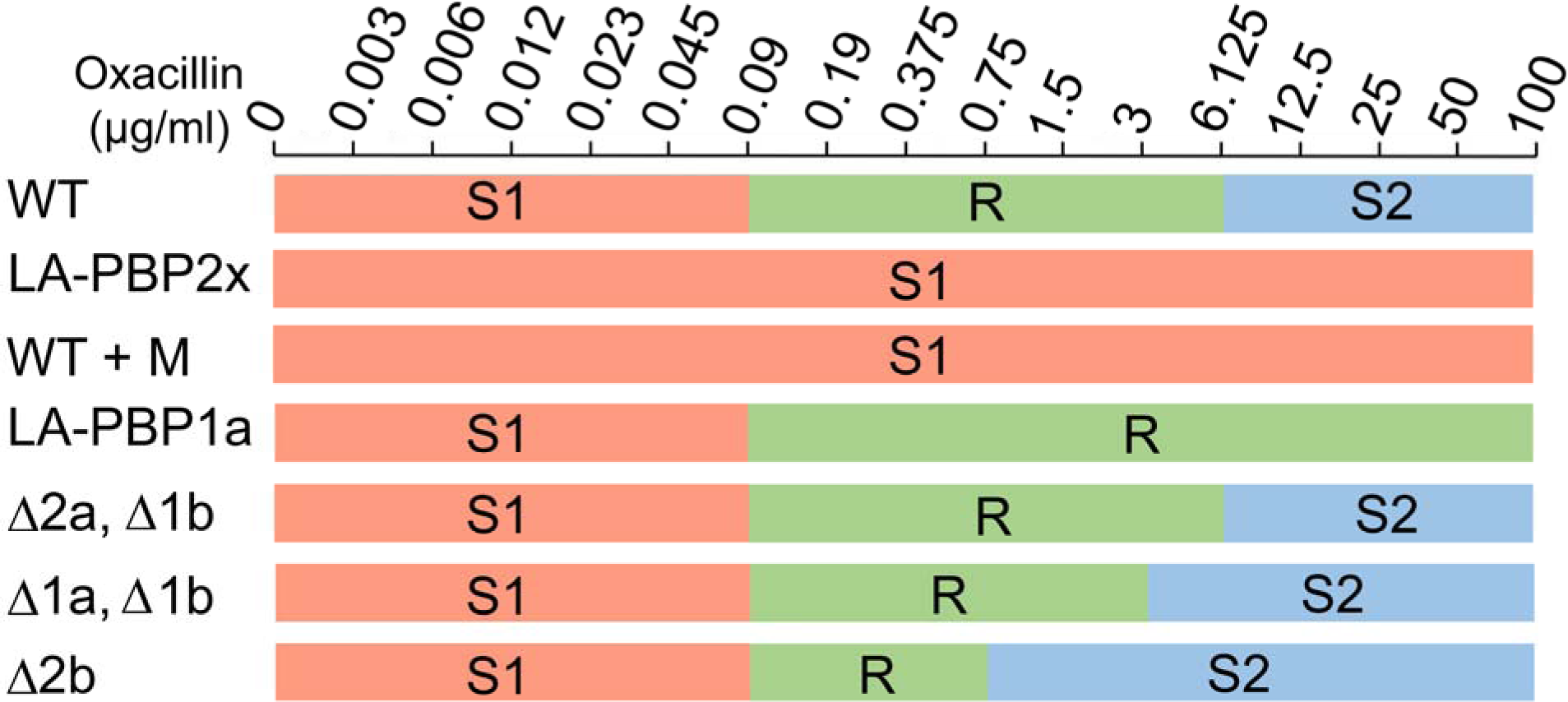
Class A PBPs are essential for transforming new peptidoglycan into a CbpD-B6-resistant form. The figure shows a schematic presentation of the sensitivity to CbpD-B6 in different genetic backgrounds based on CbpD-B6-resistance assays (see Fig. 3 and supplementary Fig. S5 for relevant growth curves). Development of the S1, R and S2 phases is indicated with th relevant concentration of oxacillin shown on top. Strain khb321 expressing a low-affinity PBP2x (LA-PBP2x) did not develop the R-phase. The same pattern was seen for WT cells (RH425) treated with moenomycin for 10 minutes (WT M) before addition of CbpD-B6. When the clas A PBP1a was changed to a low-affinity version (LA-PBP1a, strain khb332), the cells did not develop the S2-phase when treated with supra-MIC concentrations of oxacillin. The double Δ*pbp2a*/Δ*pbp1b* mutant (strain khb225) and Δ*pbp1a*/Δ*pbp1b* mutant (strain khb224) as well as a Δ*pbp2b* mutant (strain ds789) all developed the typical S1, R, S2 phases upon increasing concentrations of oxacillin. Note, the S2-phase occurred at a lower oxacillin concentration for strain khb224 and ds789 compared to wild type cells.

As three different class A PBPs are produced by *S. pneumoniae* (PBP1a, Pbp1b and PBP2a) we wondered whether or not the concerted action of all three is needed for the maturation process. To answer this question the oxacillin titration experiment was performed with a mutant strain expressing a low-affinity PBP1a protein from *S. mitis* B6. Using the same concentration range as before (0-100 µg ml^−1^ oxacillin), we only observed the S1- and R-phases in this experiment. The S2-phase had disappeared and was replaced with an extended R-phase (Fig 4 and Fig. S5B). This result shows that the activity of PBP1a alone is sufficient to transform PBP2x/FtsW-synthesized peptidoglycan into the CbpD-B6-resistant form.

The class A PBPs of *S. pneumoniae* strain R6 can be deleted one by one, and are therefore individually non-essential. PBP1a/PBP1b and PBP2a/PBP1b double mutants can also be constructed, whereas PBP1a/PBP2a double mutants are non-viable^6,7^. The fact that pneumococcal cells need either PBP1a or PBP2a to survive, indicates that these PBPs can, at least to a certain extent, substitute for each other. If the observed conversion of PBP2x/FtsW-synthesized peptidoglycan into a CbpD-B6-resistant form represents an important element in the construction of a mature pneumococcal cell wall, it would be expected that this remodelling step can be carried out also by PBP2a. To address this question, we performed the oxacillin titration experiment with a Δ*pbp2a*/Δ*pbp1b* and a Δ*pbp1a*/Δ*pbp1b* strain. In both cases we observed the usual S1, R and S2 phases (Fig. 4 and Fig. S5C and D), demonstrating that PBP2a can substitute for PBP1a in the process that transforms CbpD-B6-sensitive peptidoglycan into a resistant structure.

### PBP2b and the elongasome

Having established that class A PBPs are essential for the maturation of FtsW/PBP2x-synthesized peptidoglycan, we wanted to determine whether the maturation process also requires an active elongasome. Unfortunately, we are not aware of any β-lactam or other drug to which PBP2b is more sensitive than the other pneumococcal PBPs. Consequently, we were not able to specifically inhibit the transpeptidase activity of PBP2b without running the risk of inhibiting the activity of the other PBPs as well. Instead, we carried out the oxacillin titration experiment with a Δ*pbp2b*, Δ*lytA*, MreC^Δaa182-272^mutant strain (strain ds789), which lacks a functional elongasome. PBP2b is essential in a wild-type background, but can be deleted in a strain expressing a truncated version of the elongasome protein MreC^34^. Moreover, since pneumococci depleted in PBP2b becomes hypersensitive to LytA^8^, we deleted the *lytA* gene to avoid extensive autolysis. Deletion of *lytA* does not affect the S1-R-S2 pattern observed when wild-type pneumococci are subjected to increasing concentrations of oxacillin (Fig. S5E). When performing this experiment, we observed the usual S1-R-S2 pattern (Fig. 4 and Fig. S5F), but complete resistance was not reached when CbpD-B6 was added 10 minutes subsequent to oxacillin. However, after 15 minutes close to full resistance was obtained in cultures treated with 0.19-0.75 µg ml^−1^. This experiment shows that Class A PBPs are not dependent on a functional elongasome to convert nascent PBP2x/FtsW-synthesized peptidoglycan into a CbpD resistant form.

## Discussion

Recently it has become clear that FtsW/PBP2x and RodA/PBP2b constitute cognate pairs of interacting proteins that make up the core peptidoglycan synthesizing machineries within the pneumococcal divisome and elongasome, respectively^9,10,11^. Both couples consist of a monofunctional transglycosylase working together with a monofunctional transpeptidase. This discovery has important implications for our understanding of pneumococcal cell wall synthesis, and the role played by class A PBPs in this process. Before it was discovered that the SEDS proteins FtsW and RodA have glucosyltransferase activity, class A PBPs were considered to be the only peptidoglycan polymerases present in pneumococci. Hence, they were regarded as key components of the divisome and elongasome, and indispensable for septal as well as peripheral peptidoglycan synthesis. This way of thinking is no longer valid, and the function of class A PBPs has therefore become an open question.

Our results show that class A PBPs act downstream of the FtsW/PBP2x machinery to produce alterations in the cell wall. Class A PBPs are able to function, i.e. to convert FtsW/PBP2x-synthesized peptidoglycan into a CbpD-B6-resistant form, even when PBP2x is completely inhibited by oxacillin. Similarly, we show that class A PBPs are able to operate independently of PBP2b and the elongasome in a Δ*pbp2b* mutant. Furthermore, since the conversion process takes several minutes, the activity of class A PBPs occur subsequent to and separate in time from FtsW/PBP2x-mediated peptidoglycan synthesis. Together these findings provide three novel and important insights: i) class A PBPs have a distinct and unique role in the construction of the pneumococcal cell wall, ii) there exists a class A-mediated mechanism that remodels nascent FtsW/PBP2x-synthesized peptidoglycan into a mature CbpD-B6-resistant form, and iii) this maturation mechanism is essential.

It is well established that the divisome and elongasome constitute two separate peptidoglycan synthesizing machineries^5^. Their activities are precisely coordinated during the cell cycle, but experiments have shown that the divisome is able to operate in the absence of the elongasome and vice versa. Pneumococcal cultures treated with oxacillin (0.1 µg ml^−1^), at a concentration that inhibits PBP2x but not class A PBPs and PBP2b, give rise to highly elongated cells with no septal cross-walls (Fig. S6A and B). This demonstrates that the elongasome is active even in the absence of a functional divisome. Similar findings have been reported previously by others^5,35,36,37^. In the opposite case, several studies have shown that pneumococci are able to grow and form septal cross walls when PBP2b is depleted or deleted^8,34,38,39^. Pneumococci that are strongly depleted in PBP2b form long chains of round cells that are compressed in the direction of the long axis (Fig. S6 C and D). In the present study, we have obtained evidence that class A PBPs operate independently of the divisome and elongasome and hence function autonomously. An important question is therefore whether PBP1a, PBP2a and PBP1b operate alone or in multiprotein complexes similar to the divisome and elongasome. It has been reported that PBP1a forms a complex with CozE, MreC and MreD^40^, and that it co-immunoprecipitates with the cell cycle protein GpsB^41^. Furthermore, it has been shown that PBP2a is regulated by MacP, a substrate of the global cell cycle regulator StkP^42^. The interplay between the two PBPs and their respective partners appears to be specific, as interactions between CozE/PBP2a and MacP/PBP1a have not been detected^40,42^. Presumably, the specific partners of PBP1a and PBP2a are important for the precise spatiotemporal regulation of their activity. Together the data support a model in which PBP1a, PBP2a and PBP1b are the key players in three separate and autonomous peptidoglycan synthesizing machineries with partially overlapping functions.

The fact that class A PBP-mediated remodelling of nascent peptidoglycan is inhibited by oxacillin as well as moenomycin strongly indicates that both catalytic domains of these proteins are actively involved in the remodelling process. Hence, the remodelling mechanism most likely involves the synthesis of new glycan strands, and the incorporation of these strands into existing peptidoglycan. How could peptidoglycan synthesis by class A PBPs make the cell wall resistant to CbpD-B6? The muralytic enzyme consists of three different domains, a catalytic CHAP domain, an SH3b domain and a choline-binding domain that anchors CbpD-B6 to teichoic acid. The SH3b domain probably acts as an auxiliary module that binds peptidoglycan and facilitates the function of the catalytic CHAP domain^27^. Previous research has shown that all three domains are required for the enzyme to be active^27^. Hence, it would be sufficient to block the function of one of these domains to convert the cell wall into a CbpD-B6-resistant form. To inhibit the activity of the CHAP domain would require that the nascent peptide bridge cross-linked by PBP2x is altered to become resistant to the enzyme. A structural change in this peptide bridge might also block the binding of the SH3b domain, as the SH3b domain of lysostaphin has been reported to bind to the peptide part of the cell wall of *Staphylococcus aureus*^43^. The peptide bridges in pneumococcal peptidoglycan consists of a mixture of branched and unbranched cross-links. The branches are introduced by the aminoacyl ligases MurM and MurN. MurM catalyzes the addition of L-Ala or L-Ser, whereas the addition of the second L-Ala is catalyzed by MurN^44^. We hypothesized that if class A PBPs synthesize peptidoglycan containing only branched peptides, and the CHAP and/or SH3b domains only recognize unbranched peptidoglycan, the cell wall synthesized by class A PBPs would be resistant against CbpD-B6. This hypothesis did not receive support from the experimental data, however, as a strain lacking *murMN* behaved exactly like wild-type when subjected to the oxacillin titration assay described above (Fig. S7). Alternatively, we speculated that the SH3 domain recognizes the glycan part of the peptidoglycan instead of the peptide part. Thus, the oxacillin titration assay was performed with Δ*pgdA* and Δ*adr* mutant strains as well. The *pdgA* gene encodes a peptidoglycan *N*-acetylglucosamine deacetylase, while the *adr* gene encodes a peptidoglycan O-acetyl transferase^45,46^. The Δ*pgdA* and Δ*adr* strains displayed approximately the same S1-R-S2 pattern as the wild type strain, demonstrating that neither N-acetylation nor O-acetylation significantly affect the ability of CbpD-B6 to cleave pneumococcal peptidoglycan during the S1 and S2 phases (Fig. 8A and B). Furthermore, it is possible that class A PBP-mediated remodelling of pneumococcal peptidoglycan affects the ability of CbpD-B6 to attach to teichoic acid via its C-terminal choline-binding domain. It has been shown previously that CbpD-R6 binds to the septal region of the cell, and that the choline-binding domain alone is sufficient to target the enzyme to this localization^27^. To determine if resistance against CbpD during the R-phase is caused by reduced binding to the midcell region, we examined the binding pattern of sfGFP-CbpD-B6 to oxacillin-treated (0.8 µg ml^−1^) R-phase cells and untreated cells. The fraction of cells lacking a fluorescent signal in the midcell region was slightly higher among the R-phase cells (Fig. S4A and B), but this small difference cannot explain why R-phase cells are fully resistant against CbpD-B6. Finally, to determine whether inactivation of the class A PBPs has any effect on pneumococcal morphology, cells were treated with moenomycin for 2 hours before they were fixed and prepared for TEM. The amount of moenomycin used (0.4 µg ml^−1^) completely inhibits peptidoglycan polymerization by class A PBPs. The TEM micrographs revealed that moenomycin-treated cells had considerably thicker septal cross walls than untreated cells (Fig. S9). This observation shows that something goes wrong with the synthesis of the septal cross wall in the absence of class A PBPs. Hence, to construct a normal mature cross wall, the primary cross wall synthesised by the PBP2x/FtsW machinery must be further processed by class A PBPs. In the absence of class A PBPs, the PBP2x/FtsW machinery probably does not know when to stop and just continues to synthesise additional layers of peptidoglycan.

Considering that *S. pneumoniae* must express either PBP1a or PBP2a to be viable, class A PBPs must serve an essential function. PBP1a appears to have the most prominent role among class A PBPs, as highly β-lactam resistant pneumococci always express low-affinity versions of PBP1a in addition to PBP2x and PBP2b. We clearly show that class A PBPs together with their associated auxiliary proteins somehow remodels the primary peptidoglycan synthesized by the PBP2x/FtsW machinery. As discussed above, this remodelling might involve chemical or structural modifications of the primary peptidoglycan that inhibit the function of the CHAP, SH3b or Cbd domain of CbpD-B6. Alternatively, class A PBPs and their helper proteins might not synthesize peptidoglycan that is qualitatively different from the primary peptidoglycan synthesized by PBP2x/FtsW, but rather function as a repair machinery that mend imperfections that arise during construction and expansion of the cell wall^10^. It is conceivable that the peptidoglycan layer synthesized by PBP2x/FtsW, i.e. the divisome, is not perfect. It might not be fully homogenous but contain irregularities in the form of gaps and small holes. We speculate that CbpD-B6 use these irregularities to penetrate into and even pass through the peptidoglycan layer. This could explain the curious observation that CbpD-B6 can attack the cell wall from the inside (Fig. 2 C and D). Perhaps CbpD-B6 is not able to digest “tightly woven” peptidoglycan but depends on imperfections to get access to its substrate. We therefore propose the following scenario to explain our observations: The divisome (PBP2x/FtsW) synthesize the primary cross-wall which has structural imperfections. A few minutes later, class A PBPs and their auxiliary proteins detect and repair these imperfections making the cell wall resistant to CbpD-B6. However, since primary peptidoglycan synthesis performed by PBP2x/FtsW and Class A-mediated repair occurs simultaneously, with the latter lagging 3-4 minutes behind, there will always be some imperfect peptidoglycan in growing cells that can be attacked by CbpD. In the case where PBP2x, but not class A PBPs, is inhibited by oxacillin (phase R), all gaps and imperfections are repaired by class A PBPs before CbpD-B6 is added and the pneumococcal cells become resistant. When higher concentrations of oxacillin are used (phase S2), PBP2x and class A PBPs are inhibited at the same time, preventing repair of the most recently synthesized peptidoglycan. In sum, we believe our results are best explained by a model in which the peptidoglycan synthesizing machinery of *S. pneumoniae* consists of a divisome, an elongasome and a repairosome. The reason the repairosome consists of three different machineries, PBP1a, PBP2a and PBP1b, is probably because each have separate but overlapping functions. They might repair different types of damages and/or operate in different parts of the cell wall. Further confirmation or rejection of the repairosome hypothesis will be an important topic for future studies.

## Methods

### Cultivation and transformation of bacteria

All strains used in the present study are listed in Table S1. *E. coli* was grown in Luria Bertani broth or on LB-agar plates at 37°C containing ampicillin (100 µg ml^−1^) when necessary. Liquid cultures were grown aerobically with shaking. Chemically competent *E. coli* cells were transformed by heat-shocking at 42°C. *S. pneumoniae* was grown in liquid C medium^47^ or on Todd-Hewitt (BD Difco^®^) agar plates at 37°C. When grown on TH-agar the cells were incubated in a sealed container made anaerobically (<1% O_2_) by including AnaeroGen™ sachets from Oxoid. Transformation of *S. pneumoniae* was done by adding CSP-1 (final concentration of 250 ng ml^−1^) and the transforming DNA (50-100 ng) to one ml of exponentially growing cells at OD_550_ = 0.05. Following incubation at 37°C for two hours, transformants were selected by plating 30 µl cell culture on TH-agar plates containing the appropriate antibiotic; kanamycin (400 µg ml^−1^), streptomycin (200 µg ml^−1^) or spectinomycin (200 µg ml^−1^).

### DNA cloning

All primers used in this study are listed in Table S2. To construct pRSET-*cbpD-*B6, the *cbpD*-B6 gene from *S. mitis* B6 was amplified from genomic DNA using the primer pair so1/so2. The gene was amplified without the signal sequence encoding part, starting from codon 41. The *cbpD*-B6 amplicon was cleaved with XbaI and HindIII and ligated into pRSET A (Invitrogen) generating pRSET-*cbpD*-B6. The plasmid pRSET-sfGFP-*cbpD*-B6 was constructed by substituting the CHAP encoding part (aa 41-175) of *cbpD*-B6 with the *sf-gfp* gene. The *sf-gfp* gene was amplified using the kp116 and kp119 primers and SPH370 genomic DNA as template, and the *cbpD*-B6*-Δchap* gene was amplified from SO7 genomic DNA using the primer pair kp117/kp118. Using overlap extension PCR and the primers kp116 and kp117, *sf-gfp* was fused to *cbpD*-B6*-Δchap*. The resulting *sf-gfp-cbpD*-B6 amplicon was cleaved with NdeI and HindIII and ligated into pRSET A giving the pRSET-sfGFP-*cbpD*-B6 plasmid.

Amplicons used to transform *S. pneumoniae* were constructed by overlap extension PCR as previously described by Johnsborg et al.^22^. We employed the Janus cassette^48^ to knock out genes and to introduce recombinant DNA at desired positions in the *S. pneumoniae* genome. When substituting the native *pbp2x* gene with a low affinity version (*pbp2x*-exB6), an additional version of the native gene was ectopically expressed during transformation using the ComRS-system as described by Berg et al.^49^. The spectinomycin resistant marker *aad9* was employed to knock out *lytA* in strain ds789.

### Expression and purification of CbpD-B6

*E. coli* BL21 containing pRSET-*cbpD*-B6 was grown to OD_550_ = 0.4 – 0.5 at 37°C. Then production of CbpD-B6 was induced by adding a final concentration of 0.1 mM IPTG followed by incubation at 20°C for four hours. The cells were harvested at 5000 × g for five minutes and resuspended in 1/100 culture volume of TBS, pH 7.4. The cells were lysed using the Fast Prep method with ≤106 µm glass beads at 6.5 m s^−1^ and insoluble material were removed by centrifugation at 20 000 × g. CbpD-B6 was purified from the soluble protein fraction by performing DEAE cellulose chromatography as described by Sanchez-Puelles et al.^30^, but using TBS (pH 7.4) instead of a phosphate buffer (pH 7.0). To remove choline from the eluted CbpD-B6 protein it was dialyzed against TBS (pH 7.4) for one hour at room temperature. After concentrating the dialyzed protein to a final volume of 500 µl using an Amicon centrifugal filter (10 000 MW), it was further purified by gel filtration through a Superdex™ 75 10/300 GL column (GE healthcare) at a flow rate of 0.3 ml min^−1^ in TBS (pH 7.4).

### CbpD-B6 resistance assay

Pneumococcal cells were grown in 96-wells microtiter plates and OD_550_ was measured every five minutes. When reaching OD_550_ = 0.2, oxacillin was added in concentrations decreasing from 100 µg ml^−1^ down to 0.003 µg ml^−1^ in a two-fold dilution series. Zero antibiotic added was used as controls. In some cases, 10 µg ml^−1^ of moenomycin was added together with oxacillin. The cells were grown for 10 minutes in the presence of antibiotics before purified CbpD-B6 was added to a final concentration of five µg ml^−1^. CbpD-sensitive cells were observed as a drop in OD_550_. For the time kinetic experiments, oxacillin (0.8 µg ml^−1^) was added simultaneously to 11 parallel cell cultures grown in a 96-well microtiter plate. Then CbpD_B6_ (1 µg ml^−1^) was added to the first well at time zero, then to the second well after 1 minute and so on for 10 minutes.

### Microscopy

For TEM and SEM analysis, strain RH425 was grown to OD_550_ = 0.2 and CbpD-B6 was added to a final concentration of 0.5 µg ml^−1^. The enzyme was allowed to attack the cells for one minute at 37°C before they were fixed in a mixture of 2% (v v^−1^) formaldehyde and 2.5% (v v^−1^) glutaraldehyde. The cells were fixed on ice for one hour and then prepared for SEM and TEM imaging as previously described by Straume et al.^24^. RH425 cells grown for two hours (from OD_550_ = 0.1 to OD_550_ = 0.4) with 0.4 µg ml^−1^ moenomycin or 0.1 µg ml^−1^ oxacillin and SPH157 cells depleted for PBP2b (as described by Berg et al.^8^) was fixed and prepared for electron microscopy in the same way.

To determine the binding pattern of CbpD-B6 on sensitive and immune *S. pneumoniae* cells, a 10 ml cell culture of *S. pneumoniae* was split in two when reaching OD_550_ = 0.2. One half was left untreated, while the other half was added oxacillin to a final concentration of 0.8 µg ml^−1^. Both cultures were incubated further for 10 minutes at 37°C before formaldehyde was added to a final concentration of 2.5%. Both non-treated and oxacillin treated cells were fixed on ice for one hour. The fixed cells were washed three times in 1/5 volume of PBS, before sfGFP-CbpD-B6 (purified as described for CbpD-B6) was bound to the cell surface as described by Eldholm et al.^27^. Briefly, 100 µl of cells were applied onto a microscope glass slide (inside a hydrophobic frame made with a PAP pen) and cells were immobilized by incubation at room temperature for five minutes. Non-bound cells were rinsed off the glass by PBS. Cells were then incubated in 100 µl PBS containing 0.05% Tween 20 and 15 µg ml^−1^ sfGFP-CbpD-B6 for eight minutes at room temperature. Non-bound sfGFP-CbpD-B6 was washed off the cells by rinsing the glass slide by submerging the glass slide in five tubes each containing 40 ml PBS. Phase contrast pictures and GFP fluorescence pictures were captured using a Zeiss AxioObserver with an ORCA□Flash4.0 V2 Digital CMOS camera (Hamamatsu Photonics) through a 100x PC objective. An HPX 120 Illuminator was used as a light source for fluorescence microscopy. Images were prepared in ImageJ.

## Acknowledgments

This work was supported by grants from the Research Council of Norway (no. 240058 and 250976) and the Norwegian University of Life Sciences.

