## Supplementary material for "Class A PBPs have a distinct and unique role in the construction of the pneumococcal cell wall"

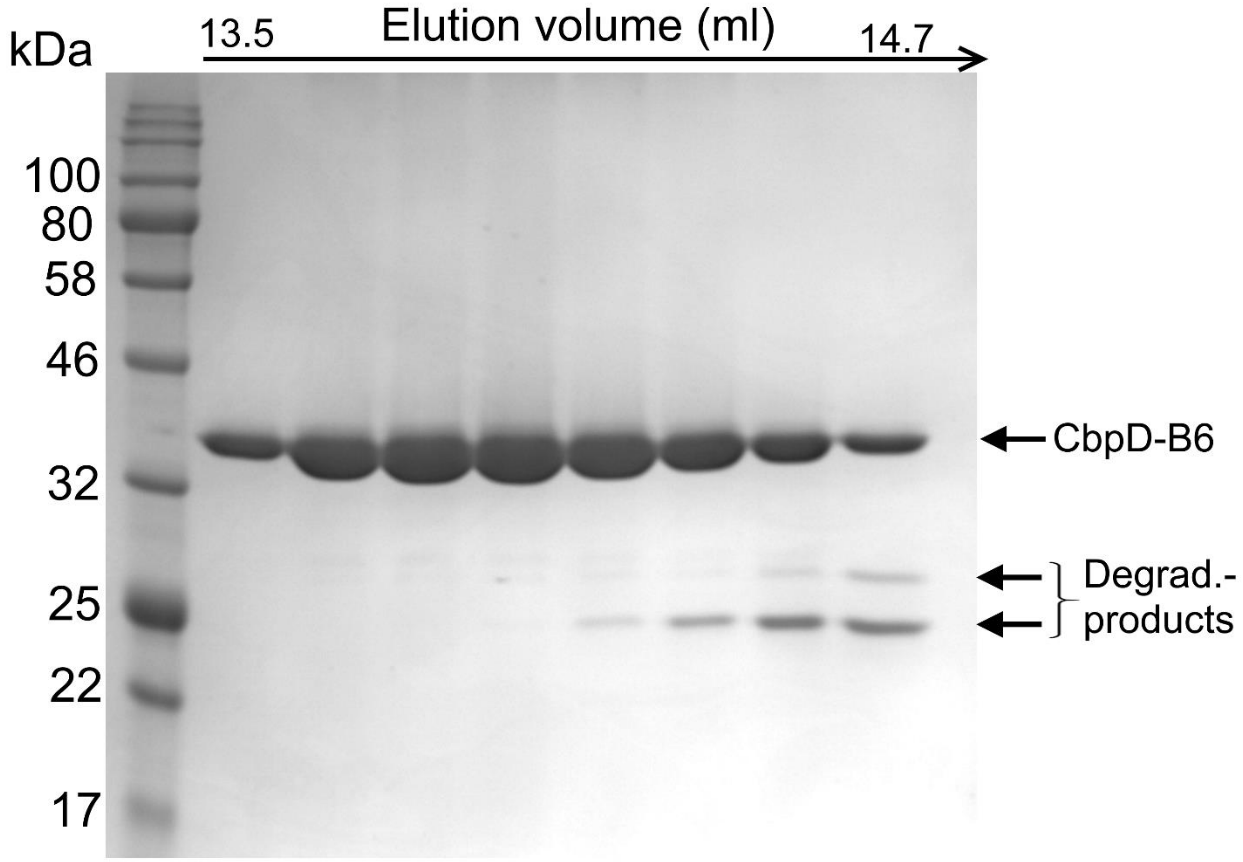


**Fig. S1.** Coomassie blue stained SDS-PAGE of CbpD-B6 purified by size exclusion chromatography (SEC).

```
CbpD-R6      MKILPFIARGTSYYLKMSVKKLVPFLVVGLMLAAGDSVYAYSRGNGSIARGDDYPAYYKN    60
CbpD-B6      MKVLPFKVTETGFSLRKSVKKVVPFLVVGLMLAASDSVYAYSGGNGSIARGDDYPAYYKN    60
             **:*** .  *.: *: ****:*************.******* ****************

CbpD-R6      GSQEIDQWRMYSRQCTSFVAFRLSNVNGFEIPAAYGNANEWGHRARREGYRVDNTPTIGS    120
CbpD-B6      GSQEIDQWRMYSRQCTSFAAFRLSNVNGFEIPRAYGNANEWGHRARREGYRVDNTPTIGS    120
             *****************.************* ****************************

CbpD-R6      ITWSTAGTYGHVAWVSNVMGDQIEIEEYNYGYTESYNKRVIKANTMTGFIHFKDLDSGSV    180
CbpD-B6      IAWSTAGTYGHVAWVSNVMGDQIEIEEYNYGYTEAYNKRIIKANTMTGFIHFKDLAGGSV    180
             *:********************************:****:*************** .***

CbpD-R6      GNSQSSASTGGTHYFKTKSAIKTEPLVSATVIDYYYPGEKVHYDQILEKDGYKWLSYTAY    240
CbpD-B6      GNSQTSASTG--------------------------------------------------    209
             ****:*****

CbpD-R6      NGSYRYVQLEAVNKNPLGNSVLSSTGGTHYFKIKSAIKTEPLVSATVIDYYYPGEKVHYD    300
CbpD-B6      --------------------------GTHYFKSKAAIKNQPLASATAIDYYYPGEKVHYD    224
                                       ****** *:***.:**.***.*************

CbpD-R6      QILEKDGYKWLSYTAYNGSRRYIQLEGVTSSQNYQNQSGNISSYGSNNSSTVGWKKINGS    360
CbpD-B6      QILEKDGYKWLSYTAYNGSRRYIQLEGVTSSQNYQNQSGNISSYGSNSSSTVGWKKINGS    284
             ***********************************************.************

CbpD-R6      WYHFKSNGSKSTGWLKDGSSWYYLKLSGEMQTGWLKENGSWYYLGSSGAMKTGWYQVSGE    420
CbpD-B6      WYHFKSNGSKSTGWLKDGSSWYYLKSSGEMQTGWLKENGSWYYLDSSGAMKTGWYQVSGK    344
             ************************* ******************.**************:

CbpD-R6      WYYSYSSGALAINTTVDGYRVNSDGERV  448
CbpD-B6      WYYSYSSGVLAVNTTVDGYRVNSDGERV  372
             ********.**:****************
```

**Fig. S2.** Amino acid sequence alignment of CbpD from *S. pneumoniae* R6 with CbpD from *S. mitis* B6. The signal sequences are shown in orange, the CHAP domains in green, SH3b domains in red and the Cbd domains in blue.

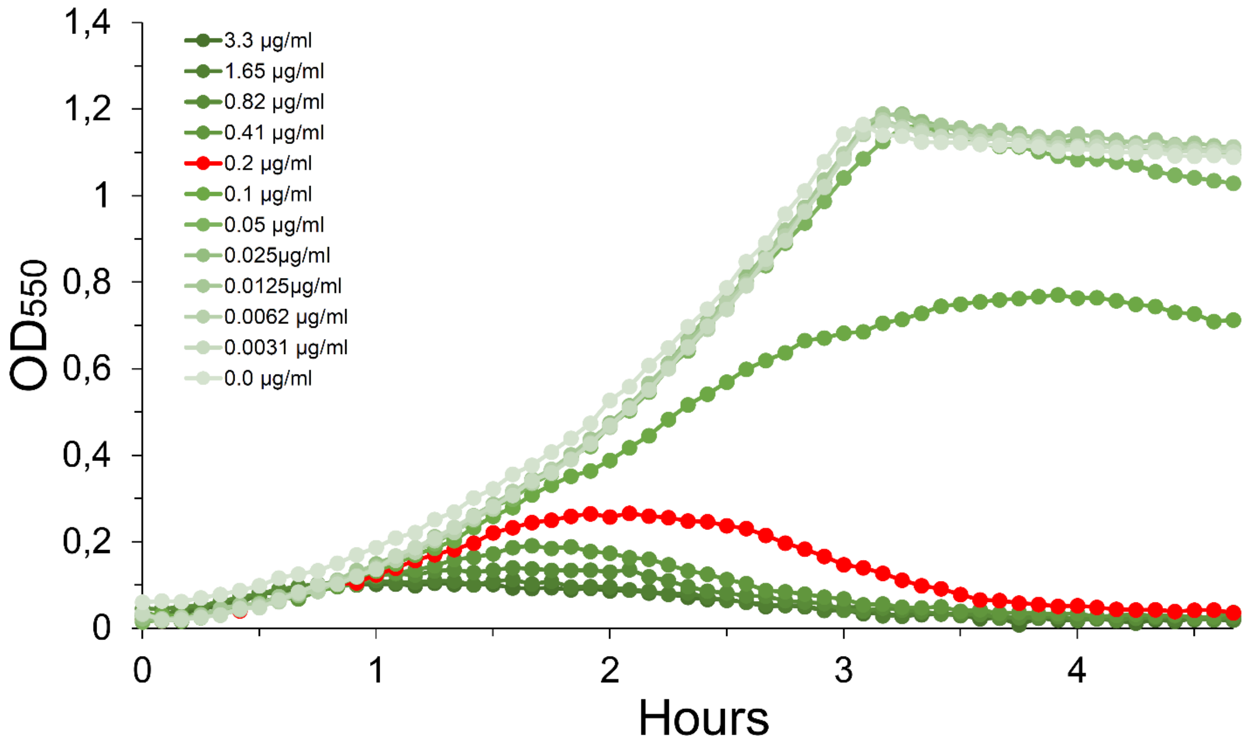


**Fig. S3.** Estimation of the oxacillin MIC value for strain RH425 in liquid culture. RH425 was grown with increasing concentrations of oxacillin in the growth medium. The concentration of the antibiotic that resulted in an approximately 80% growth retardation (red curve) compared to non-treated cells was defined as MIC. Similar results were obtained in two additional experiments.

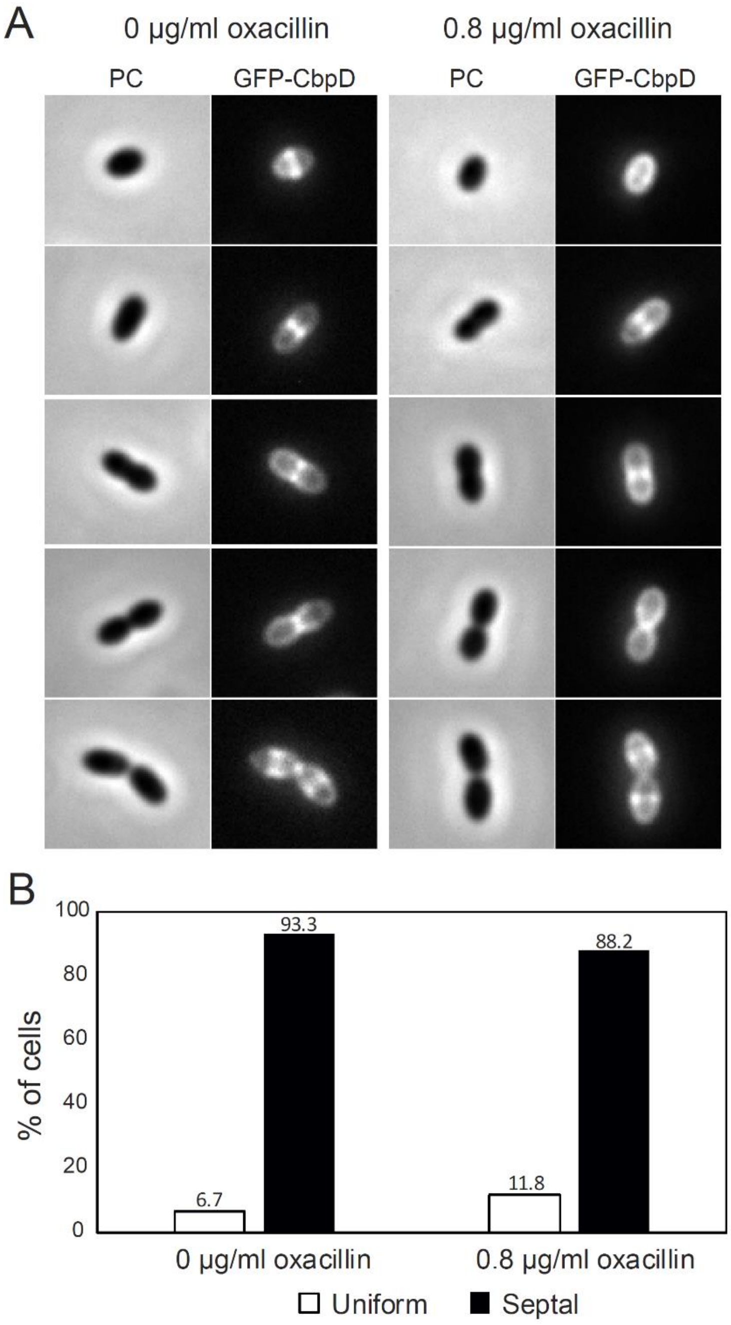


**Fig. S4.** Binding of sfGFP-CbpD-B6 to oxacillin treated *S. pneumoniae* RH425 cells. A. Binding of sfGFP-CbpD-B6 to fixed *S. pneumoniae* cells in five stages of division. Phase contrast (PC) and GFP-images of non-treated control cells (0 µg $ml^{-1}$ oxacillin, S1-phase cells) and cells treated with 0.8 µg $ml^{-1}$ oxacillin for 10 minutes (R-phase cells) are included. B. Proportion of cells with sfGFP-CbpD-B6 enriched in the septal region for both groups of cells. More than 150 cells were analyzed for each group.

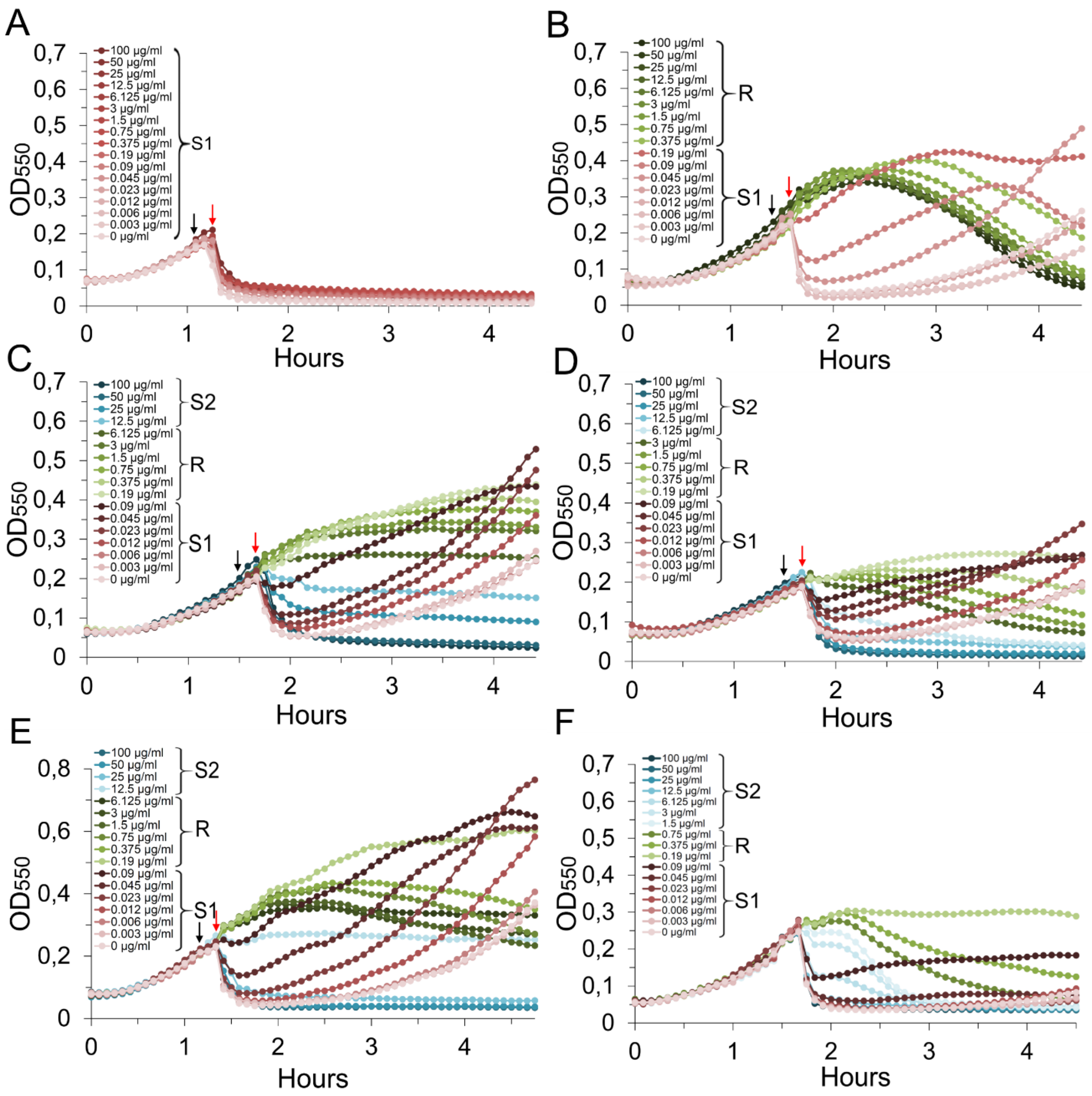


**Fig. S5.** Assay testing pneumococcal strains immunity to CbpD-B6 after treatment with different concentrations of oxacillin alone or in combination with moenomycin. The black arrows indicate when the antibiotic(s) was added, and the red arrows indicate when CbpD-B6 (5 µg $ml^{-1}$) was added. A. RH425 treated with 10 µg $ml^{-1}$ moenomycin and increasing concentrations of oxacillin, B. Low-affinity PBP1a (strain khb332), C. Strain khb225 (Δ*pbp2a*/Δ*pbp1b*), D. Strain khb224 (Δ*pbp1a*/Δ*pbp1b*), E. Strain RH14 (Δ*lytA*), F. Strain ds789 (Δ*pbp2b*, Δ*lytA*, *mreC-T*). All experiments were performed three times or more, with highly similar results.

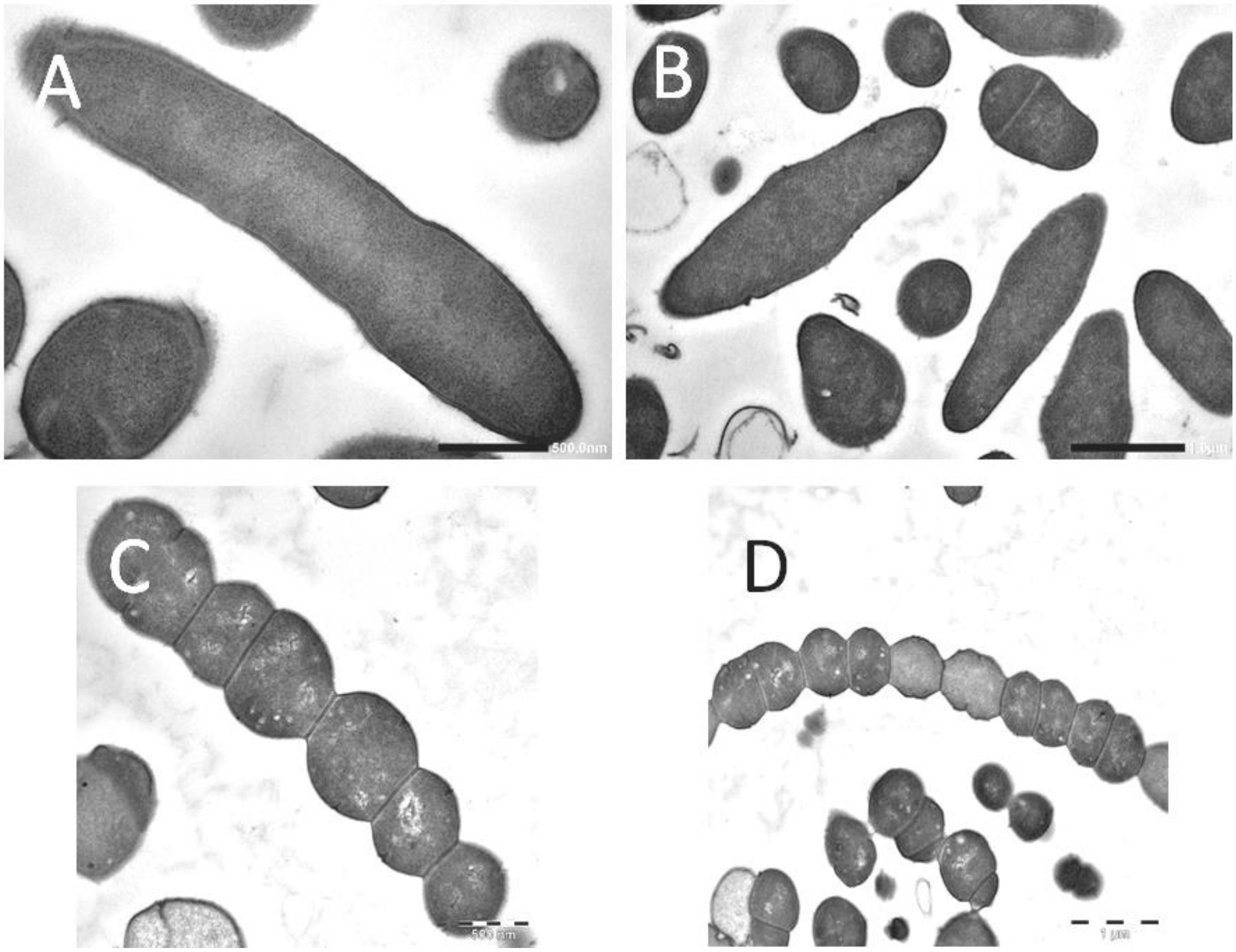


**Fig. S6.** TEM micrographs of *S. pneumoniae* RH425 cells treated with 0.1 µg ml$^{-1}$ oxacillin for 2 hours (A and B), and SPH157 cells depleted of PBP2b expression (B and C). Scale bars are 500 nm or 1 µm.

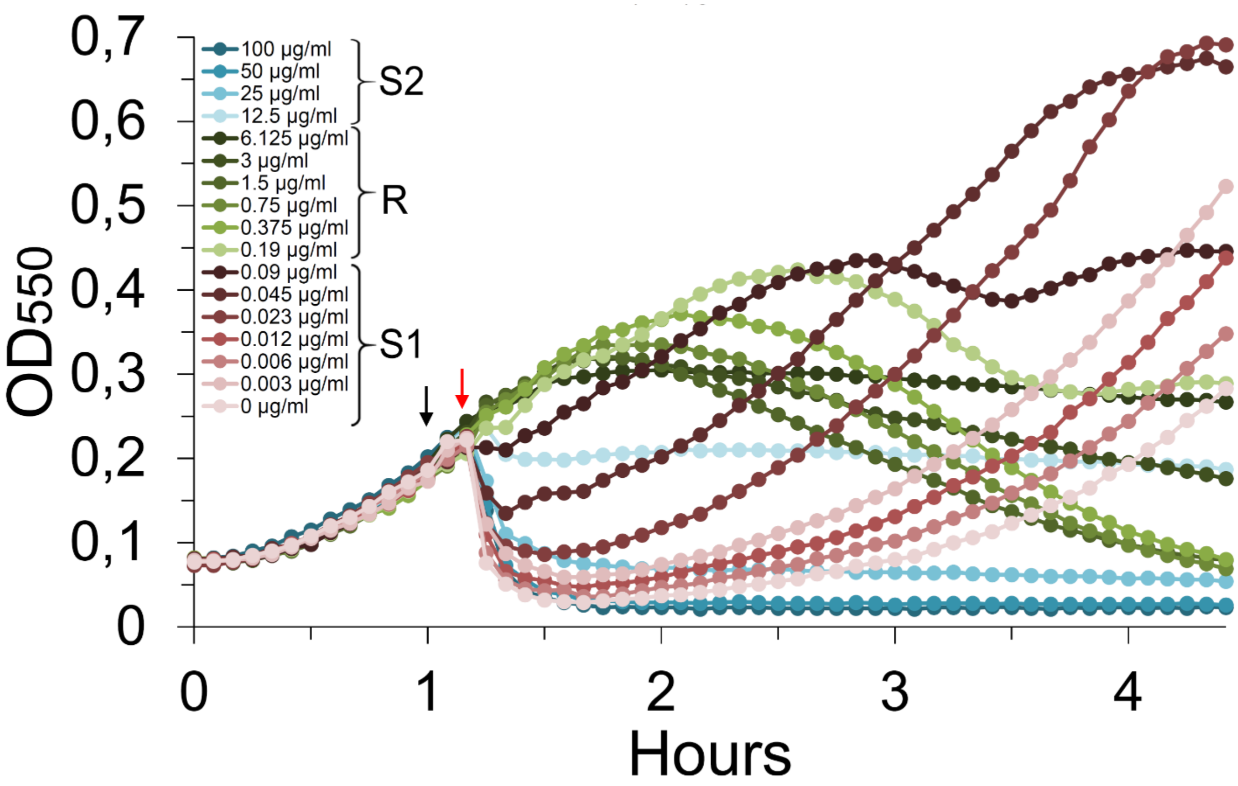


**Fig. S7.** CbpD-B6 resistance assay of strain MH110 (Δ*murMN*) after treatment with different concentrations of oxacillin added at $OD_{550}$ = 0.2 (black arrow). Addition of the enzyme (5 µg $ml^{-1}$) is indicated with a red arrow. The data presented are representative of three independent experiments.

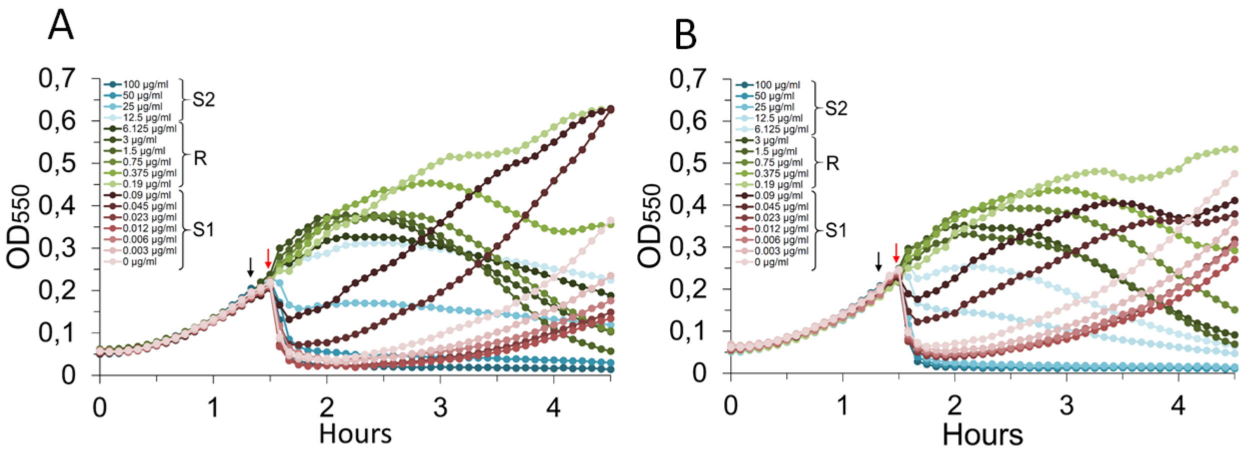


**Fig. S8.** Resistance against CbpD-B6 after oxacillin treatment of A) strain RH281 (Δ*pgdA*) and B) strain RH295 (Δ*adr*). Both mutants developed the typical S1-R-S2 phases observed for wild type *S. pneumoniae*. The Δ*pgdA* and Δ*adr* mutants were tested three times with similar results.

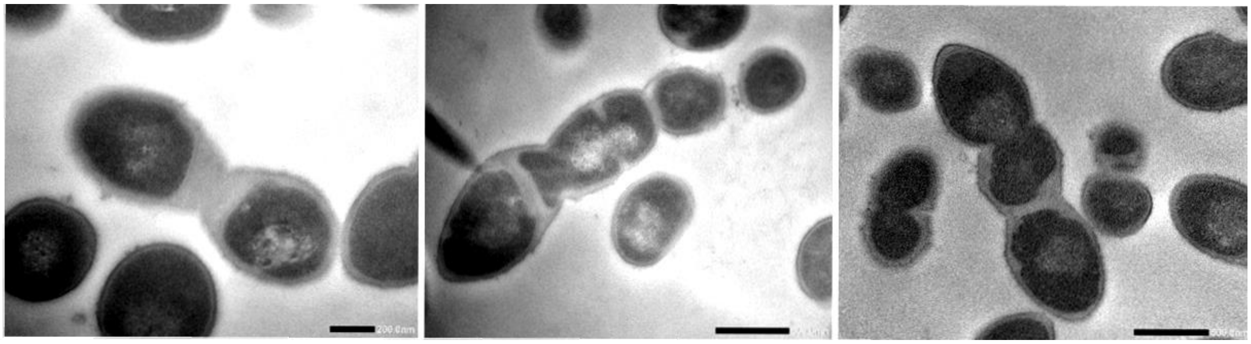

**Fig. S9.** Three representative TEM micrographs of *S. pneumoniae* RH425 cells grown with moenomycin (0.4 µg $ml^{-1}$) for 2 hours.  Thickened septal cross-walls are seen in the division zones. Scale bars are 200 nm or 500 nm.

**Table S1.** Strains used in this study.

| Strain | Relevant characteristics | Source |
|---|---|---|
| *E. coli* strains | | |
| DH5α | *E. coli* cloning host | Invitrogen |
| BL21 (DE3) | *E. coli* recombinant protein expression host | Invitrogen |
| SO3 | DH5α containing pRSET-cbpD$_{B6}$ | This study |
| SO7 | BL21 containing pRSET-cbpD$_{B6}$ | This study |
| KP5 | DH5α containing pRSET-sfGFP-cbpD$_{B6}$ | This study |
| KP6 | BL21 containing pRSET-sfGFP-cbpD$_{B6}$ | This study |
| Streptococcal strains | | |
| RH281 | Δ*comA*, Δ*comM*, Δ*pgdA*::janus; Ery$^R$, Kan$^R$ | This study |
| RH295 | Δ*comA*, Δ*adr*; Ery$^R$, Sm$^R$ | This study |
| RH425 | Δ*comA::ermAM* and streptomycin resistant; Ery$^R$, Sm$^R$ | (Johnsborg and Håvarstein, 2009) |
| RH426 | Contains the Janus cassette; Ery$^R$, Kan$^R$ | (Johnsborg and Håvarstein, 2009) |
| RH431 | Contains the Δ*lytA*::*aad9* cassette; Ery$^R$, Sm$^R$, Spc$^R$ | (Johnsborg and Håvarstein, 2009) |
| SPH131 | Δ*comA*, P1-P$_{comR}$-*comR*, P$_{comX}$-Janus; Ery$^R$, Kan$^R$ | (Berg *et al.*, 2011) |
| SPH157 | Δ*comA*, P1-P$_{comR}$-*comR*, P$_{comX}$-*pbp2b*, Δ*pbp2b*$_{wt}$; Ery$^R$, Sm$^R$ | (Berg *et al.*, 2013) |
| SPH163 | Δ*comA*, P1-P$_{comR}$-*comR*, P$_{comX}$-*pbp2x*, Δ*pbp2x*$_{wt}$::janus; Ery$^R$, Kan$^R$ | (Berg *et al.*, 2013) |
| SPH178 | Δ*comA*, P1-P$_{comR}$-*comR*, P$_{comX}$-*pbp2b*, Δ*pbp2b*$_{wt}$::janus; Ery$^R$, Kan$^R$ | (Berg *et al.*, 2013) |
| SPH370 | Δ*comA*, *sf-gfp-divIVAΔ92*; Ery$^R$, Sm$^R$ | (Straume *et al.*, 2017) |
| khb223 | Δ*comA*, Δ*pbp1b*; Ery$^R$, Sm$^R$ | This study |
| khb224 | Δ*comA*, Δ*pbp1b*, Δ*pbp1a*::janus; Ery$^R$, Kan$^R$ | This study |
| khb225 | Δ*comA*, Δ*pbp1b*, Δ*pbp2a*::janus; Ery$^R$, Kan$^R$ | This study |
| khb317 | Δ*comA*, [a]*pbp2b*$_{exB6}$, Δ(P$_{comX}$-*pbp2b*)::janus; Ery$^R$, Kan$^R$ | This study |
| khb321 | Δ*comA*, [a]*pbp2x*$_{exB6}$, Δ(P$_{comX}$-*pbp2x*)::janus; Ery$^R$, Kan$^R$ | This study |
| khb332 | Δ*comA*, [a]*pbp1a*$_{exB6}$, Ery$^R$, Kan$^R$ | This study |
| ds789 | Δ*comA*, *mreC*$^{\Delta aa183\text{-}272}$, Δ*pbp2b*, Δ*lytA::aad9;* Ery$^R$, Sm$^R$, Spc$^R$ | This study |
| MH110 | Δ*comA*, Δ*murMN*::janus; Ery$^R$, Kan$^R$ | This study |
| B6 | Penicillin resistant *S. mitis* isolated from a hospital in Bochum, Germany | (Hakenbeck *et al.*, 1998) |

[a]Extracellular part of the PBP is derived from the corresponding PBP in *S. mitis* B6.

**Table S2.** Primers used in this study.

| Primer name | Sequence 5'→3' | Source |
|---|---|---|
| Kan484.F | GTTTGATTTTTAATGGATAATGTG | (Johnsborg *et al.*, 2008) |
| RpsL41.R | CTTTCCTTATGCTTTTGGAC | (Johnsborg *et al.*, 2008) |
| **Primers used to construct Δ*pbp2a*::janus** | | |
| mts1 | GCACAACTTGTTCGTACTCTTG | This study |
| mts2 | CACATTATCCATTAAAAATCAAACGCGTTTATTTTATCATCTTCATC | This study |
| mts3 | GTCCAAAAGCATAAGGAAAGGATGCTTGTCAAAGCCTAGC | This study |
| mts4 | AGGTTTACTTCTGCAACTGTG | This study |
| **Primers used to construct Δ*pbp1a*::janus** | | |
| khb353 | GGCTTGGCTGTTATACATAAG | This study |
| khb354 | GACGGATAACCATCTCTTGAC | This study |
| mts6 | CACATTATCCATTAAAAATCAAACCTTGTTTTACCACCTAATAAATG | This study |
| mts7 | GTCCAAAAGCATAAGGAAAGCATTTATCATCCAGATTTTTCTG | This study |
| **Primers used to construct Δ*pbp1b*::janus** | | |
| mts9 | GCCTGTACTTGGTAGTTTGG | This study |
| mts10 | CATTATCCATTAAAAATCAAACGGATTTCCTCACTTTATCTATTA | This study |
| mts11 | GTCCAAAAGCATAAGGAAAGTTCTCTAAATGAAGTGGCCAATC | This study |
| mts12 | GACTATTCCAGTATAGCAC | This study |
| **Primers used to construct Δ*pbp1b*::DEL** (used in combination with mts9 and mts12) | | |
| khb276 | GTATAATAGATAAAGTGAGGAAATCCTTCTCTAAATGAAGTGGCCAATC | This Study |
| khb277 | GATTGGCCACTTCATTTAGAGAAGGATTTCCTCACTTTATCTATTATAC | This Study |
| **Primers used to construct *pbp2x*$_{exB6}$** | | |
| khb104 | GAAGTGAAGCCGATTGAGAC | (Berg *et al.*, 2013) |
| khb107 | ACACAATTCCGATAATCAAGAG | (Berg *et al.*, 2013) |
| khb339 | ACAGATTTAGCGAAGGAAGCTAAAAAGTTCACCAAACCACTCG | This study |
| khb340 | CGAGTGGTTTGGTGAACTTTTTAGCTTCCTTCGCTAAATCTGT | This study |

| | | |
|---|---|---|
| khb341 | CAGCACTGATGGAAATAAACATATTAGTCTCCTAAAG TTAATTTAATT | This study |
| khb342 | AATTAAATTAACTTTAGGAGACTAATATGTTTATTTCC ATCAGTGCTG | This study |
| **Primers used to construct $pbp2b_{exB6}$** | | |
| khb129 | CGATAAAGAAGAGCATAGGAAG | (Berg *et al.*, 2013) |
| khb132 | TCCCAATCAATGGTTTCATTGG | (Berg *et al.*, 2013) |
| ds153 | CAGACCAAGATTACAAGCAGTTCTGCTCGTGGGGAAA TTTATG | This study |
| ds154 | ACTGCTTGTAATCTTGGTCTG | This study |
| ds155 | CCAAGTATTCTGAGGGTGTGTATGCAGTCGCCCTTAA CCC | This study |
| ds156 | CACACCCTCAGAATACTTGG | This study |
| **Primers used to construct $pbp1a_{exB6}$** (used in combination with khb353 and khb354) | | |
| khb343 | GGCGGAGGAGTTTTTTTCTACTACGTCAGCAAAGCCC CAG | This study |
| khb344 | CTGGGGCTTTGCTGACGTAGTAGAAAAAAACTCCTCC GCC | This study |
| khb345 | CAGAAAAATCTGGATGATAAATGTCACTGTTGTGGTT GCTGTTG | This study |
| khb346 | CAACAGCAACCACAACAGTGACATTTATCATCCAGAT TTTTCTG | This study |
| **Primers used to amplify $cbpD_{B6}$** | | |
| so1[a] | TACGTCTAGAAATAATTTTGTTTAACTTTAAGAAGGA GATATACATATGTATTCTGGAGGAAATGGATCGATTG | This study |
| so2 | TACGAAGCTTCTATACTCGTTCTCCATCACTG | This study |
| **Primers used to construct *sf-gfp*-$CbpD_{B6}$** | | |
| kp116 | TACGCATATGAAACATCTTACCGGTTCTAAAG | This study |
| kp117 | TACGAAGCTTCTATACTCGTTCTCCATCACTG | This study |
| kp118 | CTAGTGGAGCGGCCGCAGGTGGTGGTGGTGGTGCTGG TGGCAGTGTTGGG | This study |
| kp119 | CCCAACACTGCCACCAGCACCACCACCACCACCTGCG GCCGCTCCACTAG | This study |
| **Primers used to amplify Δ*lytA*::*aad9*** | | |
| VE17 | TGTATCTATCGGCAGTGTGAT | (Eldholm *et al.*, 2009) |
| VE20 | TCAACCATCCTATACAGTGAA | (Eldholm *et al.*, 2009) |
| **Primers used to amplify Δ*murMN::janus*** | | |
| VE47 | ACCAGTAGTCATGGAAGCAAA | (Berg *et al.*, 2011) |
| VE50 | CACATTATCCATTAAAAATCAAACTTCCTACTCTCTTT CCTCCA | (Berg *et al.*, 2011) |

| | | |
|---|---|---|
| khb198 | CTAAACGTCCAAAAGCATAAGGAAAGGATGAAAAAG TCAGTATTTAGATT | (Berg *et al.*, 2011) |
| khb199 | CACAATTTCAGACACCAGAGC | (Berg *et al.*, 2011) |

[a]restriction sites are underlined
